# Reconstructing cell cycle and disease progression using deep learning

**DOI:** 10.1101/081364

**Authors:** Philipp Eulenberg, Niklas Köhler, Thomas Blasi, Andrew Filby, Anne E. Carpenter, Paul Rees, Fabian J. Theis, F. Alexander Wolf

**Affiliations:** Helmholtz Zentrum München – German Research Center for Environmental Health, Institute of Computational Biology, Neuherberg, Munich, Germany; Department of Physics, Arnold Sommerfeld Center for Theoretical Physics, LMU München, Munich, Germany; Imaging Platform at the Broad Institute of Harvard and Massachusetts Institute of Technology, Cambridge, Massachusetts, USA; Flow Cytometry Core Facility, Faculty of Medical Sciences, Newcastle University, Newcastle upon Tyne, UK; College of Engineering, Swansea University, Singleton Park, Swansea, UK; Department of Mathematics, TU München, Munich, Germany

## Abstract

We show that deep convolutional neural networks combined with non-linear dimension reduction enable reconstructing biological processes based on raw image data. We demonstrate this by reconstructing the cell cycle of Jurkat cells and disease progression in diabetic retinopathy. In further analysis of Jurkat cells, we detect and separate a subpopulation of dead cells in an unsupervised manner and, in classifying discrete cell cycle stages, we reach a 6-fold reduction in error rate compared to a recent approach based on boosting on image features. In contrast to previous methods, deep learning based predictions are fast enough for on-the-fly analysis in an imaging flow cytometer.

## Introduction

A major challenge and opportunity in biology is interpreting the increasing amount of information-rich and high-throughput single-cell data. Here, we focus on imaging data from fluorescence microscopy (Pepperkok and Ellenberg, 2006), in particular from imaging flow cytometry (IFC), which combines the fluorescence sensitivity and high-throughput capabilities of flow cytometry with single-cell imaging (Basiji *et al.*, 2007). Imaging flow cytometry is unusually well-suited to deep learning as it provides very high sample numbers and image data from several channels, that is, high-dimensional, spatially correlated data. Deep learning is therefore capable of processing the dramatic increase in information content — compared to spatially integrated fluorescence intensity measurements as in conventional flow cytometry (Brown and Wittwer, 2000) — in IFC data. Also, IFC provides one image for each single cell, and hence does not require whole-image segmentation.

Deep learning enables improved data analysis for high-throughput microscopy as compared to traditional machine learning methods (Eliceiri *et al.*, 2012; Blasi *et al.*, 2016; Jones *et al.*, 2009; Dao *et al.*, 2016). This is mainly due to three general advantages of deep learning over traditional machine learning: there is no need for cumbersome preprocessing and manual feature definition, prediction accuracy is improved, and learned features can be visualized to uncover their biological meaning. In particular, we demonstrate that this enables reconstructing continuous biological processes, which has stimulated much research effort in the past years (Gut *et al.*, 2015; Bendall *et al.*, 2014; Trapnell *et al.*, 2014; Haghverdi *et al.*, 2016). Only one of the other recent works on deep learning in high-throughput microscopy discusses the visualization of network features (Pärnamaa and Parts, 2016), but none deal with continuous biological processes (Chen *et al.,* 2016a; Kraus *et al.,* 2016; Dürr and Sick, 2016; Kandaswamy *et al.,* 2016; Pärnamaa and Parts, 2016).

When aiming at an understanding of a specific biological process, one often only has coarse-grained labels for a few qualitative stages, for instance, cell cycle or disease stages. While a continuous label could be efficiently used in a regression based approach, qualitative labels are better used in a classification-based approach. In particular, if the ordering of the categorical labels at hand is not known, a regression based approach will fail. Also, the detailed quantitative information necessary for a continuous label is usually only available if a phenomenon is already understood on a molecular level and markers that quantitatively characterize the phenomenon are available. While this is possible for cell cycle when carrying out elaborate experiments where such markers are measured (Gut *et al.,* 2015; Blasi *et al.,* 2016), in many other cases, this is too tedious, has severe side effects with unwanted influences on the phenomenon itself or is simply not possible as markers for a specific phenomenon are not known. Therefore, we propose a general workflow that uses a deep convolutional neural network combined with classification and visualization based on non-linear dimension reduction (Fig. 1).

**Figure 1.**
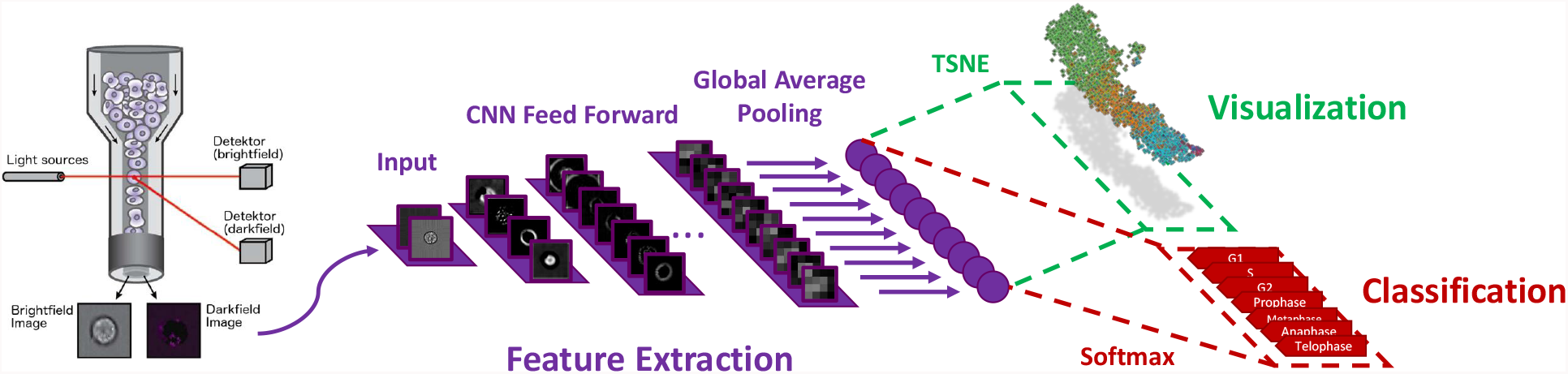
Overview of analysis workflow. Images from all channels of a high-throughput microscope are uniformly resized and directly fed into the neural network, which is trained using categorical labels. The learned features are used for both classification and visualization.

## Materials and Methods

The primary dataset in this paper consists in raw IFC images of 32,266 asynchronously growing immortalized human T lymphocyte cells (Jurkat cells), which was previously analyzed using traditional machine learning (Blasi *et al.,* 2016; Hennig *et al.,* 2016). Images of these cells can be classified into seven different stages of cell cycle (Figure 2), including phases of interphase (G1, S and G2) and phases of mitosis (Prophase, Anaphase, Metaphase and Telophase). In this data set, ground truth is based on the inclusion of two fluorescent stains: propidium iodine (PI) to quantify each cell’s DNA content and the mitotic protein monoclonal #2 (MPM2) antibody to identify cells in mitotic phases. These stains allow each cell to be labeled through a combination of algorithmic segmentation, morphology analysis of the fluorescence channels, and user inspection (Blasi *et al.,* 2016). Note that 97.78% of samples in the dataset belong to one of the interphase classes G1, Sand G2. The strong class imbalance in the dataset is related to the fact that interphase lasts — when considering the actual length of the biological process — a much longer period of time than mitosis.

**Figure 2.**
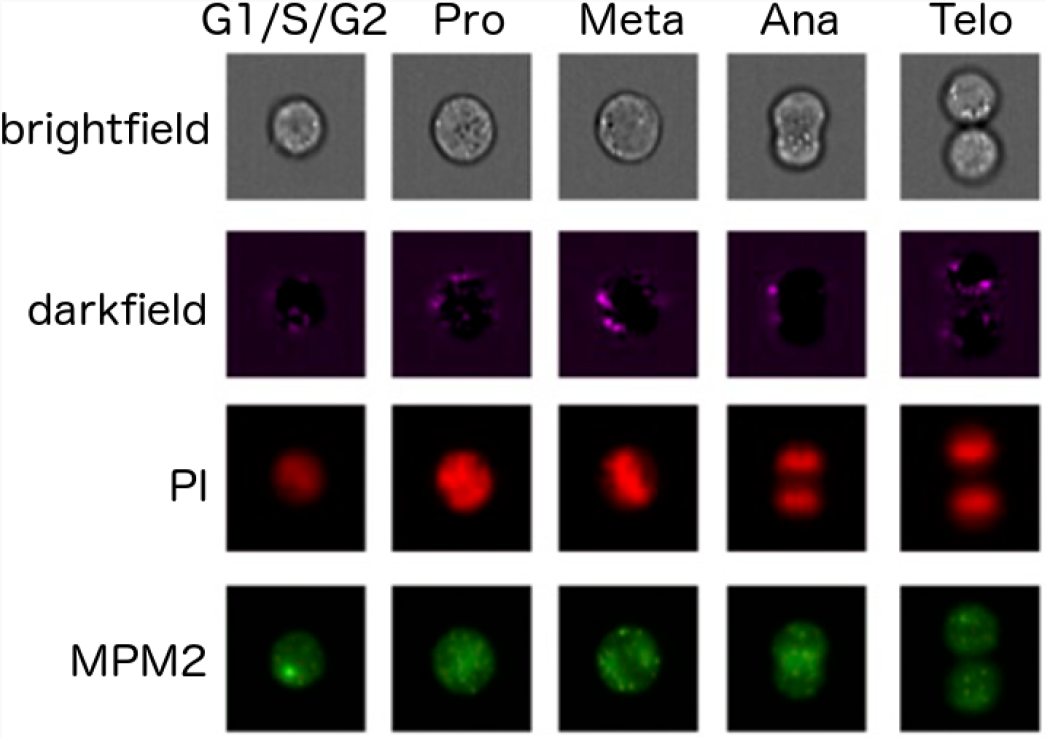
Representative images for the cell cycle stages as measured in brightfield, darkfield and fluorescence channels. Seven cell cycle stages define seven classes. We only show one representative image for the interphase classes G1, S, and G2, which can hardly be distinguished by eye.

To substantiate the generality of our results, we study a second dataset that was collected with a technology other than IFC and is related to a biological process other than cell cycle: 30.000 publicly available images from the Diabetic Retinopathy Detection Challenge (2015). Diabetic retinopathy is the leading cause of blindness in the working-age population of the developed world. It is diagnosed by trained humans based on the presence of lesions visible in color fundus photographies of the retina and is classified into four disease states: “healthy”, “mild”, “medium’’, and “severe”.

Recent advances in deep learning have shown that deep neural networks are able to learn powerful feature representations (Krizhevsky *et al.,* 2012; Vincent *et al.,* 2010; Szegedy *et al.,* 2015; LeCun *et al.*, 2015). We adapt the widely used “Inception” architecture (Szegedy *et al.*, 2015) and, for the IFC data, optimize it for treating the relatively small input dimensions. The architecture consists in 13 three-layer “dual-path” modules (Suppl. Fig. S3), which process and aggregate visual information at an increasing scale. These 39 layers are followed by a standard convolution layer, a fully connected layer and the softmax classifier. Training this 42-layer deep network does not present any computational difficulty, as the first three layers consist in reduction dual-path modules (Suppl. Fig. S3b), which strongly reduce the original input dimensions prior to convolutions in the following normal dual-path modules. The number of kernels used in each layer increases towards the end, until 336 feature maps with size 8 × 8 are obtained. A final average pooling operation melts the local resolution of these maps and generates the last 336-dimensional layer, which serves as an input for both classification and visualization.

This neural network operates directly on uniformly resized images. It is trained with labeled images using stochastic gradient descent with standard parameters (see Suppl. Notes). For the IFC data, we focus on the case in which only brightfield and darkfield channels are used as input for the network, during training, visualization and prediction. As stated before, this case is of high interest as a fluorescent markers might affect the biological process under study or adequate markers are not known. We note, however, that technical imperfections in the IFC data capture might always lead to a minor amount of fluorescence signal, activated by a fluorescence channel, in the darkfield and brightfield channels, a phenomenon known as “bleed through” (see Suppl. Notes).

## Results

To show how learned features of the neural network can be used to visualize, organize and biologically interpret single-cell data, we study the activations in the last layer of the neural network (Donahue *et al.*, 2013). We refer to this as studying the *activation space representation* of the data. The approach is motivated by the fact that the neural network strives to organize data in the last layer in a linearly separable way, given that it is directly followed by a softmax classifier. Distances from the separating hyperplanes in this space can be interpreted as similarities between cells in terms of the features extracted by the network. Cells with similar feature representations are close to each other and cells with different class assignments are far away from each other. As we will see, this gives a much more fine-grained notion of biological similarity than provided by the class labels used for labeling the training set. Evidently, it automatically generalizes to the unseen, new data in the validation data set.

The activation space of our network’s last layer has 336 dimensions and is much too high-dimensional to be accessible for human interpretation. We use non-linear dimension reduction to visualize the data in a lower dimensional space, in particular, t-distributed stochastic neighbor embedding (tSNE) (van der Maaten and Hinton, 2008) (see Suppl. Video).

### Reconstructing cell cycle progression

In this visualization, we observe that the Jurkat cell data is organized in a long stretched cylinder along which cell cycle phases are ordered in the chronologically correct order (Fig. 3a). This is remarkable as the network has been provided with neither structure within the class labels nor the relation among classes. The learned features evidently allow reconstructing the continuous temporal progression from the raw IFC data, and by that define a continuous distance between the phenotypes of different cell cycle phases.

**Figure 3.**
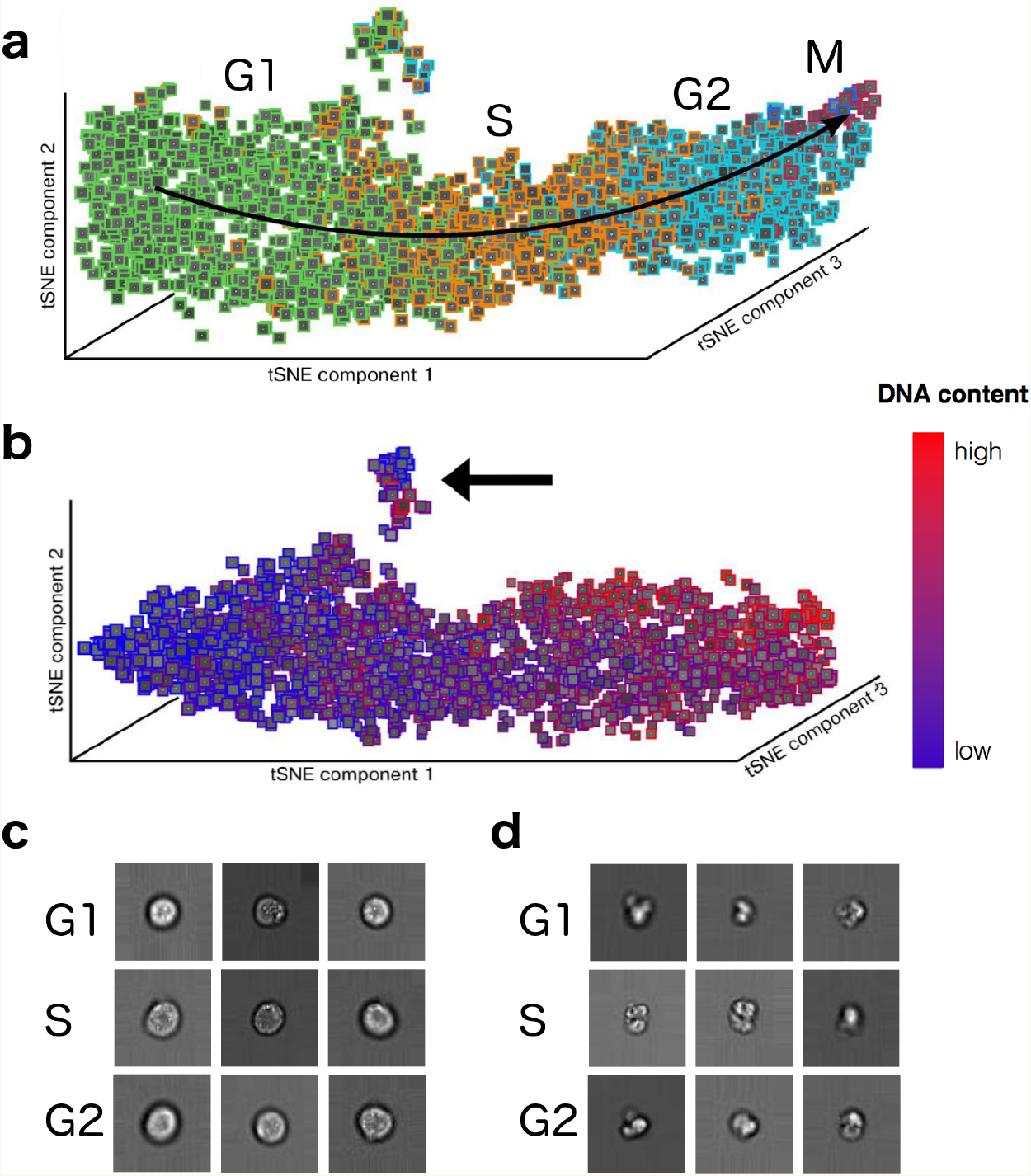
Cell-cycle reconstruction and detection of abnormal cells. **a,** tSNE visualization of the validation dataset in activation space representation. All interphase classes (G1, S, G2) and the two mitotic phases with the highest number of representatives are shown (Prophase: red, Metaphase: blue). Telophase and Anaphase are not visible due to their low number representatives. **b,** tSNE visualization of data from the interphase classes (G1, S, G2) in activation space. The color map now shows the DNA content of cells. A cluster of damaged cells is indicated with an arrow. **c,** Randomly picked representatives from the bulk of undamaged cells. **d,** Randomly picked representatives from the cluster of damaged cells.

We separately visualized just those cells annotated as being in the interphase classes (G1, S, G2) (Figure 3b) and colored them with the DNA content obtained from one of the fluorescent channels (PI). The DNA content reflects the continuous progression of cells in G1, S and G2 on a more fine-grained level. Its correspondence with the longitudinal direction of the cylinder found by tSNE demonstrates that the temporal order learned by the neural network is accurate even beyond the categorical class labels.

### Detecting abnormal cells in an unsupervised manner

Both tSNE visualizations (Fig. 3a,b) produce a small, separate cluster highlighted with an arrow in Fig. 3b. This cluster is learned in an unsupervised way as cell cycle phase labels provide no information about it: it contains cells from all three interphase classes. While cells in the bulk have high circularity and well defined borders (Fig. 3c), cells in the small cluster are characterized by morphological abnormalities such as broken cell walls and outgrowths, signifying dead cells (Fig. 3d).

### Deep learning automatically performs segmentation

We interpret the data representation encoded in one of the trained intermediate layers of the neural network by inspecting its activation patterns using exemplary input data from several classes (Fig. 4). These activation patterns are the essential information transmitted through the network. They show the response of various kernels on their input. By inspecting the activation patterns, we obtain an insight into what the network is “focusing on” in order to organize data. We observe a strong response to features that arise from the cell border thickness (Fig. 4, map 1), to area-based features (Fig. 4, map 2), as well as cross-channel features. For example, map 4 in Fig. 4 shows high response to the difference of information from the brightfield channel, as seen in map 2, and scatter intensities, as seen in map 3. A strong response of the neural network to area-based features as in map 2 could indicate that the network learned to perform a segmentation task.

**Figure 4.**
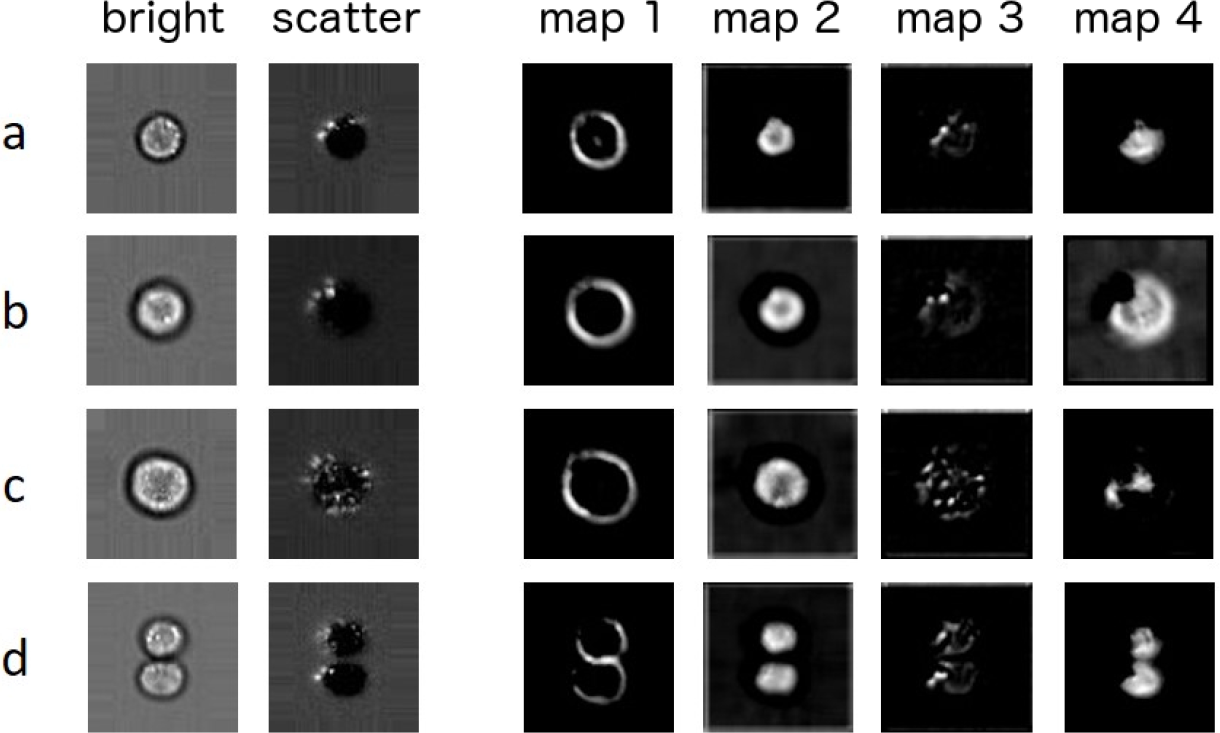
Exemplary activation patterns of intermediate layers. Plotted are activations after the second convolutional module for examples of single cells from four different phases: **a,** G1 **b,** G2 **c,** Anaphase and **d,** Telophase. The response maps mark regions of high activation. Map 1 responds to the cell boundaries. Map 2 responds to the internal area of the cells. Map 3 extracts the localized scatter intensities. Map 4 constitutes a cross-channel feature, which correlates with the difference of map 2 and 3.

### Deep learning outperforms boosting for cell cycle classification

We study the classification performance of Deep learning on the validation data set shown in Fig. 3. We first focus on the case in which G1, S and G2 phases are considered as a single class. Using five-fold cross-validation on the 32,266 cells, we obtain an accuracy of 98.73%±0.16%. This means a 6-fold improvement in error rate over the 92.35% accuracy for the same task on the same data in prior work using boosting on features extracted via image analysis (Blasi *et al.*, 2016). The confusion matrix obtained using boosting show high true positive rates for the mitotic phases (Fig. 5a). For example, no cells in Anaphase and Telophase are wrongly classified, as indicated by the zeros in the off-diagonal entries of the two lower rows of the matrix (Fig. 5a). This means high *sensitivity*, most cells from mitotic phases are correctly classified as such. Still this comes at the price of low *precision*: many cells from the interphase class are classified as mitotic phases, as indicated by the high numbers in the off-diagonal entries of the first row of the matrix (Fig. 5a). Deep learning, by contrast, achieves high sensitivity and precision, leading to an almost diagonal confusion matrix (Fig. 5b). Further Deep learning allows to classify all seven cell cycle stages with an accuracy of 79.40%±0.77% (see Suppl. Notes and Suppl. Fig. S2).

**Figure 5.**
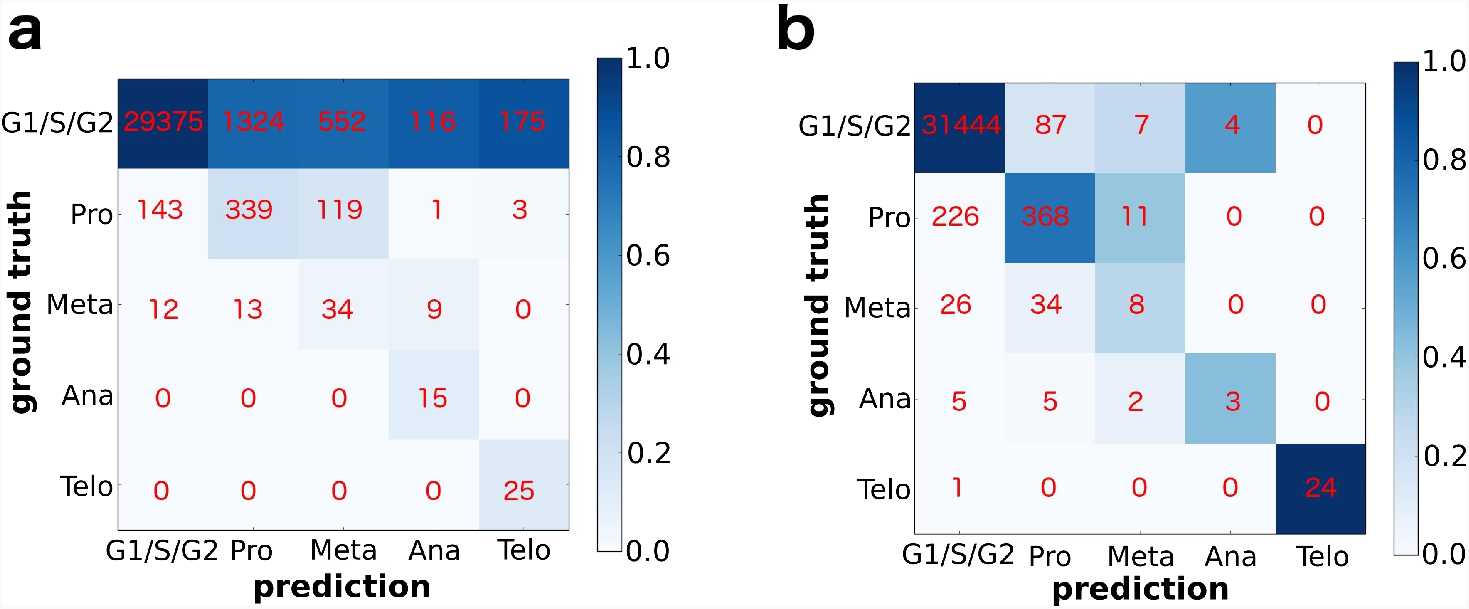
Confusion matrices for boosting and Deep learning for classification of five classes. To compare with previous work (Blasi *et al.*, 2016), the three interphase phases (G1, S, G2) are treated as a single class. Red numbers denote absolute numbers of cells in each entry of the confusion matrix, that is, diagonal elements correspond to precision. Coloring of the matrix is obtained by normalizing absolute numbers to column sums. **a,** Boosting (Blasi *et al.*, 2016), which leads to 92.35% accuracy. **b,** Deep learning, which leads to 98.73%±0.16% accuracy.

### Reconstructing disease progression

Consider now the second dataset, which deals with diabetic retinopathy (DR), as described above. Having trained the neural network with four different qualitative disease states, “healthy”, “mild”, “medium”, and “severe”, we observe a reconstructed disease progression (Fig. 6) for 8000 samples in the validation dataset, that is, the four disease states are ordered along disease severity, even though the network has not been provided with the ordering information. Similar to the cell cycle example, the ordering ensures that only neighboring classes overlap, as visible from the tSNE plot (Fig. 6a). Hence, the confusion matrix (not shown) displays a similar close-to tridiagonal structure as for the cell cycle (Fig. 5b and Suppl. Fig. S2a).

**Figure 6.**
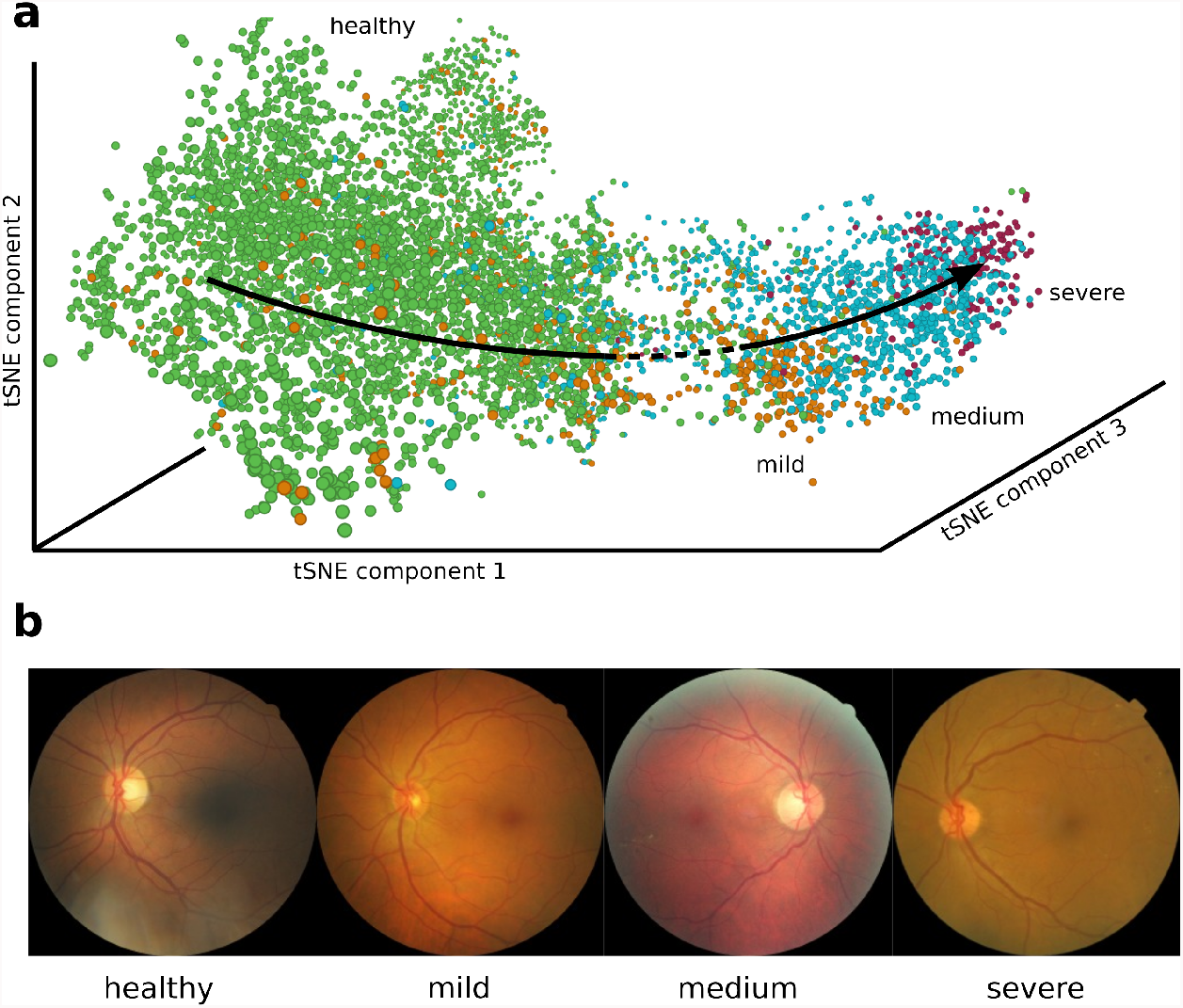
Reconstruction of disease progression in diabetic retinopathy. **a,** tSNE visualization of activation space representation, colored according to the disease states. **b,** Randomly chosen images for each class.

## Discussion

The visualization of the data as encoded in the last layer of the network using tSNE demonstrates how deep learning overcomes a well known issue of traditional machine learning. When trained on a continuous biological process using discrete class labels, traditional machine learning often fail to resolve the continuum (Eliceiri *et al.*, 2012). Reconstructing continuous biological processes though is possible in the context of so-called pseudotime algorithms (Bendall *et al.*, 2014; Trapnell *et al.*, 2014; Haghverdi *et al.*, 2016). For the cell-cycle it has been demonstrated by Gut *et al.* (2015), but in a very different setting. These authors measured five stains that uniquely define the cell cycle and then applied a pseudotime algorithm (Bendall *et al.*, 2014) within this five-dimensional space. This procedure is only possible if stains that correlate with a given process of interest are known, if they do not interact with the process and if the elaborate experiments for measuring the intensity of these stains can be carried out. We, by contrast, use raw images directly and the learned features of the neural network automatically constitute a feature space in which data is continuously organized. In the Suppl. Notes, we demonstrate that pseudotime algorithms fail at solving this much harder problem.

Deep learning is able to reconstruct continuous processes based on categorical labels as adjacent classes are morphologically more similar than classes that are temporally further separated. If this assumption does not hold, also pseudotime algorithms fail to reconstruct a process. This can be better understood when inspecting Fig. 6a, where we show the tSNE visualization of the validation set for the diabetic retinopathy (DR) data. Samples are organized in the correct order of progression through disease states, from healthy to severe DR. However, between the healthy cluster (green) and the mild DR cluster (orange), one observes an area of slightly reduced sampling density (dashed line). This should not be attributed to “less data points having been sampled in this region” but should be seen as a consequence of the fact that the overlap between the “healthy” stage and the “mild” stage is smaller than the overlap of the diseased stages among each other. If there was no overlap between “healthy” and “mild” stages, the tSNE would show a complete separation of the healthy cluster from the rest of the data. Such a behavior is typically observed if the underlying data is not sampled from a continuous process.

The unsupervised detection of a discrete cluster of abnormal cells for the Jurkat cell data indicates that the neural network learns the cluster of abnormal cells independently of the cell-cycle-label based training. The model is therefore not only capable of resolving a biological process, but generates features that are general enough to separate incorrectly labeled cells that do not belong to the process. None of the mentioned pseudotime algorithms is capable of this. This shows the ability of deep learning to find unknown phenotypes and processes without knowledge about features or labels. Also, there is a high practical use of the detection of damaged cells. The Jurkat cell data set has been preprocessed using the IDEAS^®^ analysis software to remove images of abnormal cells. In particular, out of focus cells were removed by gating for images with gradient RMS and debris was removed by gating for circular objects with a large area. The discovery of a cluster of abnormal cells shows the limitations of this approach and provides a solution to it.

An advantage of using a neural network for cell classification in IFC is its speed. Traditional techniques rely on image segmentation and measurement, time-consuming processes limited to roughly 10 cells per second. Neural network predictions, by contrast, are extremely fast, as the main computation consists in parallelizable matrix multiplications (“forward propagations”), which can be performed using optimized numeric libraries. This yields a roughly 100-fold improvement in speed to about 1000 cells per second with a single GPU. Aside from much faster analysis of large cell populations, this opens the door to “sorting on the fly”: imaging flow cytometers currently do not allow physically sorting individual cells into separate receptacles based on measured parameters, due to these speed limitations.

## Conclusion

Given the compelling performance on reconstructing the cell cycle, we expect deep learning to be helpful for understanding a wide variety of biological processes involving continuous morphology changes. Examples include developmental stages of organisms and the progression of healthy states to disease states, situations that have often been non-ideally reduced to binary classification problems. Ignoring intrinsic heterogeneity has likely hindered a deeper insight into the mechanisms at work. Analysis as demonstrated here could reveal morphological signatures at much earlier stages than previously recognized.

Our results indicate that reconstructing biological process is possible for a wide variety of image data, if enough samples are available. Although generally lower-throughput in terms of the number of cell processed, conventional microscopy is nevertheless still high-throughput and can usually provide higher resolution images than IFC. Furthermore, given that multi-spectral methods are advancing rapidly, imaging mass spectrometry is allowing dozens of labeled channels to be acquired (Bodenmiller *et al.*, 2012; Angelo *et al.*, 2014). Due to its basic structure and high flexibility, our deep learning framework can accommodate a large increase in the number of available channels.

## Acknowledgments

F.A.W. acknowledges support by the Helmholtz Postdoc Programme, Initiative and Networking Fund of the Helmholtz Association. P.R. and A.E.C. acknowledge the support of the Biotechnology and Biological Sciences Research Council/ National Science Foundation under grant BB/N005163/1 and NSF DBI 1458626.

## Supplemental Notes

### Pseudotime-based reconstruction of cell cycle

To compare our deep learning results with the class of pseudotime algorithms (Bendall *et al.*, 2014; Trapnell *et al.*, 2014; Haghverdi *et al.*, 2016; Gut *et al.*, 2015), we use Diffusion Pseudotime (DPT) (Haghverdi *et al.*, 2016) in the implementation of Scanpy (Wolf *et al.*, 2017). DPT has recently been very favorably discussed by the authors of Monocle (Qiu *et al.*, 2017), one of the most established pseudotime algorithms (Trapnell *et al.*, 2014), and is more robust than Wanderlust (Bendall *et al.*, 2014), the underlying algorithm of Gut *et al.* (2015). Using DPT, we infer the progression of Jurkat cells based on different sets of features:

- deep learning (this work),
- classical image features (Blasi *et al.*, 2016),
- marker intensity (Gut *et al.*, 2015).

For this, we focus on predicting the DNA content of cells, which measures the progression of cell cycle during G1, S and G2 phase as in Fig. 3b. Deep learning outperforms classical image extraction techniques (Fig. S1a versus b): Both in the tSNE and the pseudotime versus DNA content scatter, a high correlation is only visible in the case of deep learning (Pearson correlation of 0.56 versus 0.021). Note that the tSNE in Fig. S1a shows the same data as Fig. 3b, but in a two-dimensional tSNE.

**Supplementary Figure S1.**
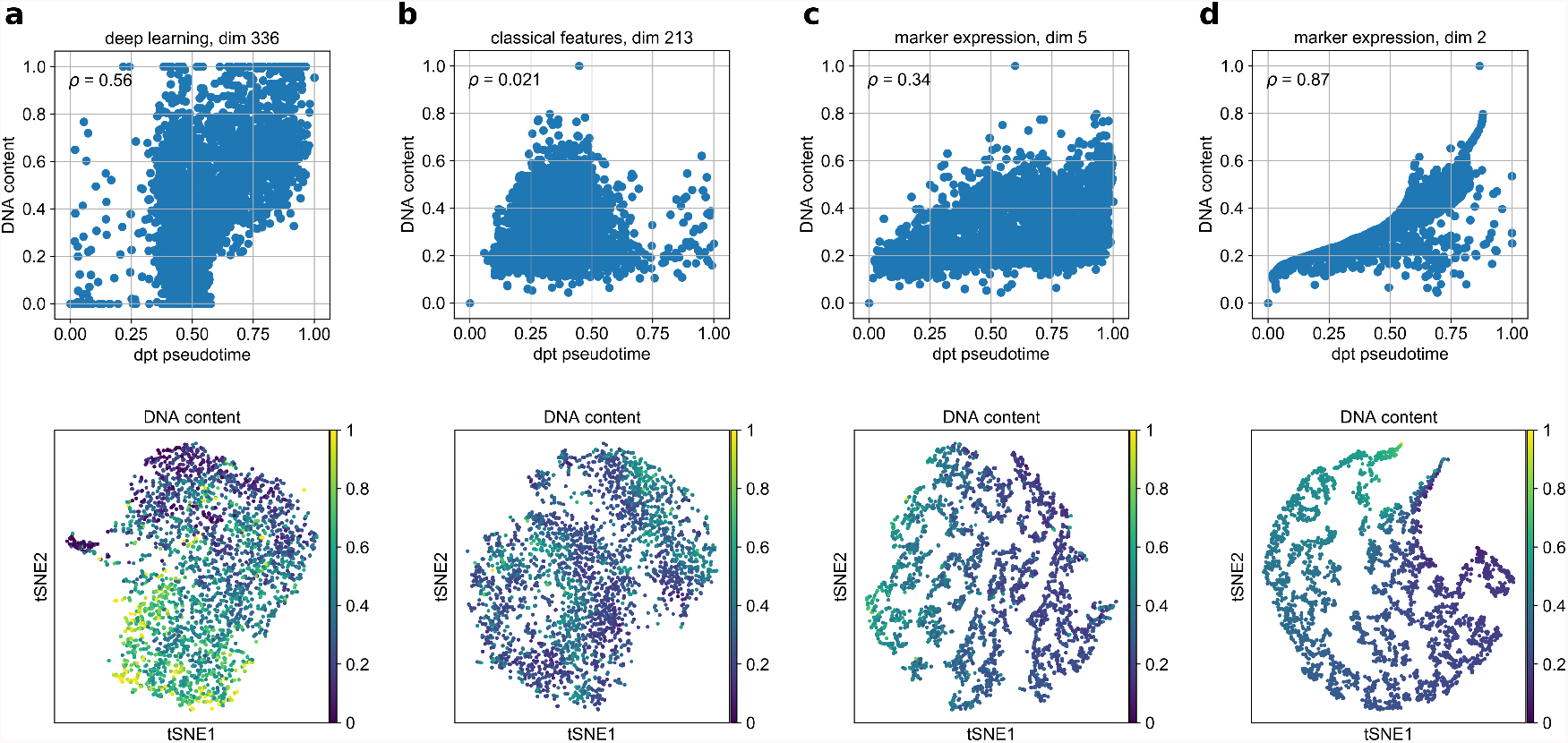
Comparison of pseudotime inference based on different feature sets. The first row shows scatter plots of DNA content versus pseudotime. The second row shows tSNE visualizations of the data represented in the respective features spaces. **a,** Deep learning (this work), **b,** classical features (Blasi *et al.*, 2016) **c,** marker intensity, analogous to Gut *et al.* (2015), with a feature space of dimension 5, but here, only one dimension stores the marker that measures DNA content. **d,** Same as **c**, but for a feature space of dimension 2.

When using the marker expression directly as an input for the pseudotime reconstruction, as done by Gut *et al.* (2015), the quality of the reconstruction depends on the number of informative features. Here, we consider the five-dimensional (Fig. S1c) and the two-dimensional case (Fig. S1d). In both cases, the first feature is the marker intensity measuring DNA content, and other features are non-specific classical image features. It is not astonishing that in the latter case (Fig. S1d), pseudotime correlates very well with the DNA content. If, as done by Gut *et al.* (2015), only the informative feature is used as an input, one obtains perfect agreement. It is important to note that this requires knowledge of the markers and measuring the known markers, both of which poses strong constraints on the problem of interest. Also, note that only the deep learning based features are not only able to reconstruct the process, but at the same time separate the cluster of abnormal cells (tSNE of Fig. S1a, clearly marked in the 3-dimensional version Fig. 3b).

Finally, note that the marker expression cannot only be used in pseudotime algorithms but also directly as an input for a regression based machine learning evalution. This has been studied for the cell-cycling Jurkat cells by Blasi *et al.* (2016), who in addition to the classification discussed in the main text above, trained a regression boosting on the classical image features using the marker for DNA content as a continuous label. Not astonishingly, the results for this were much better (Pearson correlations of 0.786 to 0.894). The disadvantages of such an analysis and the advantages of our approach have already been discussed.

More visualizations and details on the analysis of this section are available here.

### Classification of all seven cell-cycle phases

We also evaluated the full seven-class problem in which the three interphase classes are considered individually. Here, we obtain an accuracy of 79.40%±0.77%. This number serves as an orientation for what deep learning could be able to achieve on this particularly hard classification problem — G1, S and G2 are extremely difficult to distinguish (see Fig. 2), even when using information from the fluorescence channels. The accuracy might therefore be affected by wrong labelling, it might be higher if all fluorescence channels were used as input for the neural network, and it might be slightly lower if “bleed through” enriched brightfield and darkfield images. If high classification accuracy is of importance, and one is not only interested in visualizing and interpreting the data, these questions have to be answered from case to case. Their answer depends in particular on how labels are generated and how many channels of the IFC are used.

**Supplementary Figure S2.**
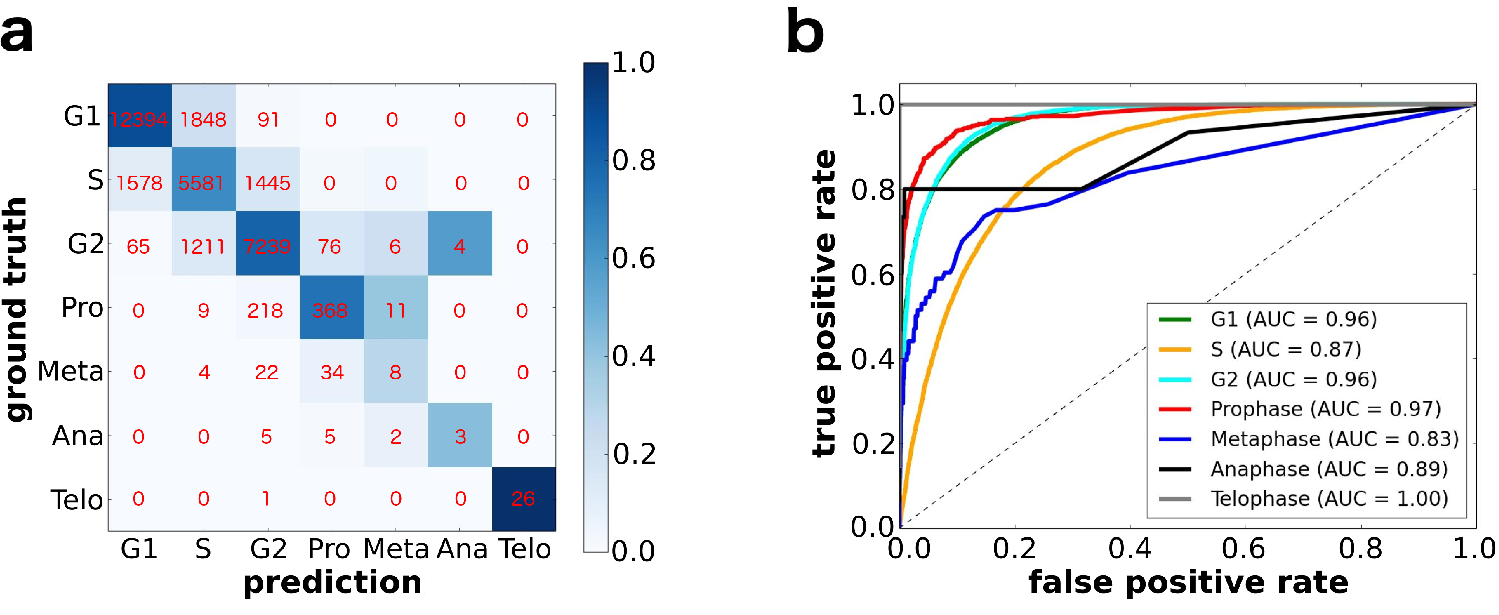
Performance of Deep learning for classification of seven classes. **a,** Confusion matrix. Red numbers denote absolute numbers of cells in each entry of the confusion matrix. Coloring of the matrix is obtained by normalizing absolute numbers to column sums, that is, diagonal elements correspond to precision. **b,** Class-specific Receiver Operating Characteristics.

Here, we confirm that the considerably lower accuracy as compared to the five-class problem results primarily from cells in the S phase being wrongly classified as either G1 or G2 (Fig. S2a). This is also shown by the Receiver Operating Characteristic, which relates the true positive rate (sensitivity) with the false positive rate (fall-out) as the classification threshold changes (Fig. S2b). Integrating the curve to obtain the standard performance metric “Area under the curve” (AUC). Even though the AUC for the S phase is still high with 0.87, it is the lowest among the majority classes (G1, S, G2), and therefore has a strong effect on the accuracy. Overall we find that all seven classes yield high values, greater than 0.85, and four of the seven classes, yield very high values, greater than 0.95.

### Technical Notes

#### Preprocessing

Our algorithmic workflow of cell cycle analysis with Deep Neural Networks begins with brightfield and darkfield images from the cells. In order to allow uniform training of our network on the whole dataset, we resize the images to 66 × 66 pixels by stretching the border pixels. We choose this method over individual image rescaling to avoid the destruction of possibly important size relation information between cells.

The data set used in this paper has been preprocessed using the IDEAS^®^ software (Merck Millipore Inc.) to remove images of abnormal cells. In particular, we removed out of focus cells by gating for images with gradient RMS and removed debris by gating for circular cells with a large area.

#### Network architecture

Figure S3 shows normal and reduction dual-path modules, the basic elements of the deep learning architecture discussed in the main text. Kernel sizes, stride and number of filters are indicated in the figure.

The network architecture consists of 42 layers, which results in a total number of parameters of about 2 mio. It is build up starting with 3 dual-path reduction modules, followed by 10 normal dual-path modules, one pooling layer, one fully connected layer and the softmax layer. Each dual-path module consists in 3 layers: a convolution layer, a batch normalization layer and an activation layer. Although there is no “big” fundamental difference between dualpath and standard convolution modules, dual-path based networks tend to converge a little better in practice, since the gradient flow from pooling and convolution in the reduction module counteracts the vanishing gradient problem: not the entire gradient gets multiplied by approx 10*−*4 convolutional weight, pooling just lets it through.

**Supplementary Figure S3.**
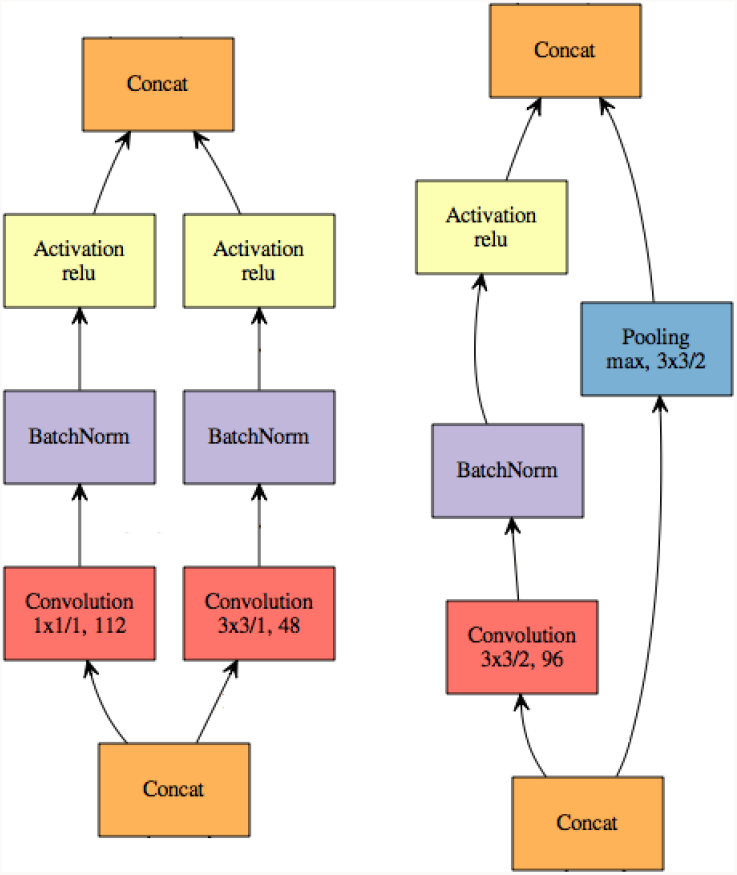
Dual-path modules. **a,** Normal dual-path module. **b,** Reduction dual-path module. The numbers beneath the convolution operations indicate the kernel sizes, stride and the number of filters.

In the first (input) layer, all IFC channels are combined in a linear operation by feeding them in the channel — which equals the color — dimension of the convolution input. This means the convolution uses kernels which convolve over all channels simultaneously. The number of 3×3 kernel weights then is nine times the number of channels. Increasing the number of channels simply increases the “kernel depth” in the color dimension, and hence, is trivial.

We note that the specific choice of the architecture is a matter of experience and cannot be further justified. Readers might use these parameters to reproduce our results or produce similar results on similar problems. We also note that there is some freedom in the specific choice of the architecture, small modifications will not qualitatively alter the results.

#### Training details

The network was trained for 100 epochs using stochastic gradient descent with standard parameters: 0.9 momentum, a fixed learning rate of 0.01 up to epoch 85 and of 0.001 afterwards as well as a slightly regularizing weight decay of 0.0005. Training took around 7 h and was stopped manually by inspecting convergence cross-entropy.

#### Implementation

For the results presented in this paper, we implemented neural networks using the MxNet framework (Chen *et al.*, 2016b) on a NVIDIA Titan X GPU. MxNet is lightweight, fast and memory efficient and available from https://github.com/dmlc/mxnet. Due to the fast progress in the development of deep learning software packages, in the meanwhile, we have also implemented and successfully tested our architecture using TensorFlow, which is available from https://github.com/tensorflow/tensorflow. The user might choose the software package according to personal preferences.

#### Nonlinear dimension reduction

We use tSNE (van der Maaten and Hinton, 2008). See Scanpy (Wolf *et al.*, 2017) for a package that provides several non-linear dimension reduction methods for visualizing single-cell data.

#### Bleed through

The data acquired using the ImageStream was fully compensated using typical control images (see Ref. (Filby *et al.*, 2011)) so the image tiffs would have minimal bleed through between channels. We could not detect even a slight indication of bleed through in the Jurkat cell data, neither upon inspection by eye, nor upon correlating the integrated intensity of each fluorescence channel with the integrated intensity of bright and darkfield channels, respectively. We then checked the existence of bleed through in the Cytometer used for data generation by switching off the light source of the brightfield channel, while keeping the fluorescence excitation on. We would then expect zero intensity in the brightfield images, but instead measured a slight intensity stemming from the fluorescence channels. This common technical aspect of IFC measurements merits an own investigation and will appear elsewhere. Here, our aim is to compare methodologies rather than to claim absolute levels of accuracy.

